# The Decision of the Optimal Rank of a Non-negative Matrix Factorization Model for Gene Expression Datasets Utilizing Unit Invariant Knee Method

**DOI:** 10.1101/2022.04.14.488288

**Authors:** Emine Güven

## Abstract

**Background:** There is a great need to develop a computational approach to analyze and exploit the information contained in gene expression data. Recent utilization of non-negative matrix factorization (NMF) in computational biology has served its capability to derive essential details from a high amount of data in particular gene expression microarrays.

**Objective:** A common problem in NMF is finding the proper number rank (r) of factors. Thus, various techniques have been suggested to select the optimal value of rank factorization (r).

**Method:** This study focused on the unit invariant knee (UIK) method to calculate factorization rank (basis vector) of the non-negative matrix factorization (NMF) of gene expression data sets is employed. Because the UIK method requires an extremum distance estimator (EDE) that is eventually employed for inflection and identification of a knee point, this study finds the first inflection point of curvature of RSS of the proposed algorithms using the UIK method on gene expression datasets as a target matrix.

**Results:** Computation was conducted for the UIK task using the esGolub data set of R studio, and consequently, the distinct results of NMF was subjected to compare on different algorithms. The proposed UIK method is easy to perform, free of a priori rank value input, and does not require initial parameters that significantly influence the model’s functionality.

**Conclusion:** This study demonstrates that the UIK method provides a credible prediction for both gene expression data and precisely estimating of simulated mutational processes data with known dimensions.

## 1 INTRODUCTION

Nonnegative matrix factorization (NMF) algorithms have been advanced for the application fields of bioinformatics, artificial intelligence (AI), signal processing systems, and music signal processing systems (1). The NMF puzzle with a parts-based illustrated an algorithm has been formulated to solve the problem by Lee and Seung (2). Furthermore, various algorithms have been established to develop a solution to the NMF problem depending on the field (3–6).

Several approaches have been developed for clustering samples, mutational processes, and genes expression levels that draw similar expression motifs (2, 7–9). Nevertheless, these approaches have severe constraints to catch the entire framework essential in the data. Moreover, they generally highlight the dominant forms in a data set and collapse to detect different signatures and etiquette. Thus, we need an unbiased technique for deciding many clusters without eye involvement and capable of utilizing a computational program.

A common problem in multivariate data analysis methods such as Factor Analysis (FA), Principal Component Analysis (PCA), Cluster Analysis (CA), and Non-negative Matrix Factorization (NMF) is to detect the proper number r of factors, principal components, clusters, and ranks, respectively. This study aims to utilize the “Unit Invariant Knee” (UIK) method for obtaining related biological and molecular correlations in gene expression data. The UIK method is used to catch compositions essential for the data and, by systematizing both the features and patients, to offer biological understanding. The approach is based on a knee point and its unit invariant estimation using Extremum Distance Estimator (EDE) method introduced by Christopoulous et al., 2012 (10). In this regard, NMF decomposes gene expression data set into fragments of evocative features such as metagenes, signatures, and mutational signatures. In contrast, this method was stated that applying conventional factorization techniques, such as principal component analysis (PCA) or factor analysis (FA), to World Values Survey Wave 5 data for United States data (11) reveals factors (elements) do clearly explain questionnaire (1 = ‘Not at all like me,.., 6 = ‘Very much like me’) interpretation (12,13).

Here, given an NMF method and a data set (a target matrix), the tens of thousands of genes regarding a small number of signatures. Samples can then be studied, drawing their gene expression patterns about expression motifs of the signatures. The signatures define an interesting decomposition of genes, analogous to motifs of Hutchins et al., 2008 study of choosing the first value where the RSS curvature presents an inflection point. To detect this inflection point, in other words, signatures or metagenes, we use the machinery of the UIK method working on data sets. Furthermore, the signature expression motifs define a robust clustering of samples. This project synthesized the UIK technique for model selection that uses alternative parsing and evaluates its robustness. Finally, this study applied the combination of NMF and the UIK method to simplify cancer classification tasks by clustering tumor samples. As a result, we can illustrate numerous sturdy decompositions of leukemia sets in experimental and simulated data.

## 2. MATERIALS AND METHOD

### 2.1. Non-negative Matrix Factorization

Given a target matrix V^m×n^, non-negative matrix factorization (NMF) identifies non-negative matrices such that N^m×r^ and M^r×n^ (i.e., with all entries ≥ 0) to present the matrix decomposition as:

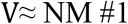

In practice, this study is typically viewed N as a basis or metagenes matrix, and mixture coefficient matrix and metagene expression profiles to refer to matrix N. The rank factorization is chosen such that r ≤ min(m, n). The goal behind this selection is to explain and split the details classified among V into r factors: the columns of N. Given a matrix V^m×n^, NMF fisnds two non-negative matrices *N*^*m*×*r*^ and M^r×n^ (i.e., with all elements ≥ 0) to represent the decomposed matrix as:

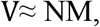

for instance, by natural demanding of non-negative N and M to minimize the reconstruction error:

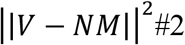

It is considered a gene expression data set characterized by the expression levels of m genes (probes) by n samples of unique tissues, cells, cell lines, time points, or experiments. The number m of genes is usually from hundreds to thousands, and the n of experiments or patients is usually 100 for gene expression research. The gene expression data set are presented by a matrix of expression V of size NxM, whose rows consist of m genes expression levels and columns of n samples.

The aim is to identify a small number of rank factorization, each defined as a positive linear combination of the V target matrix. The positive linear combination of metagenesis is described by the gene expression motif of the samples. To obtain a dimensional reduction of the microarray data and evaluate the distinctions among samples, non-negative matrix factorization (NMF) was implemented utilizing R statistical environment version 3.6.3 with the “NMF” package (14).

### 2.2. Cross Validation

This study used cross-validation to select an optimal number of implicit elements in non-negative matrix factorization. The goal of NMF is to obtain low dimensional N and M with all non-negative elements by minimizing reconstruction error | |**V** − **NM**| |^**2**^. Leaving out a single entry of V, e.g. **V**_**ab**_, and implement NMF of the resulting matrix may produce differently than the actual result. In other words, finding N and M minimizing reconstruction error over all non-missing entries:

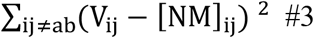

Consequently, it can be predicted the left-out element *V*_*ab*_ by calculating [WH]_ab_ and then calculate the prediction error as:

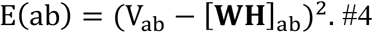

One can repeat this process by crossing out all entries of *V*_*ab*_ one at a time and adding up the error of prediction over all, *a*_*a*_ and *a*_*b*_. This will lead to the predicted residual sum of squares (PRESS) value. The PRESS value is defined as *E*(r) = ∑_ab_ E(ab), which will strongly depend on the rank of r. The prediction error, E(r) will have a minimum defined as an ‘optimal rank’ r.

Since the NMF has to be reiterated for each crossed out value and might also be difficult to code (depending on the target matrix entries and how smooth it is to implement NMF with missing values), this can be a computationally expensive procedure. For instance, in PCA, one can avoid this by crossing out entire rows of V, which eventually speeds up the computing (15). All the traditional cross-validation rules can apply here. Therefore, not including multiple entries instead of a single entry and iterating the computation process by bootstrapping the entries instead of looping over all the entries, both techniques can help speed up the procedure.

### 2.3. Proposed UIK method

Hutchins et al., 2008 previously demonstrated how the variation in the RSS of the estimated matrix resulting from the NMF analysis reveals a robust approximation of the proper number of elements (r) (8). They employed Lee and Seung’s algorithm to select r, in which the plot of the RSS presents the first inflection point. In practice, the rank factorization r can be computed with a considerably smaller number of iterations, typically twenty to thirty runs for each value of r. In contrast, an optimal NMF interpretation requires a couple of hundred random re-starts, computationally expensive.

As Figure 1 shows, this study aims to find an inflection where r meets the proper number of the factorization rank. The utilization of Unit Invariant Knee (UIK) methodology for the identification of knee (elbow) point of a curve is immensely advantageous in a wide variety of studies constantly to locate the optimal number of ‘components’ on a scree plot of PCA or FA In many cases, utilization is uik(x,y), where x is the vector of ranks, components, clusters, or factors and y is the related vector of Residual Sum of Squares (RSS).

**Figure 1.**
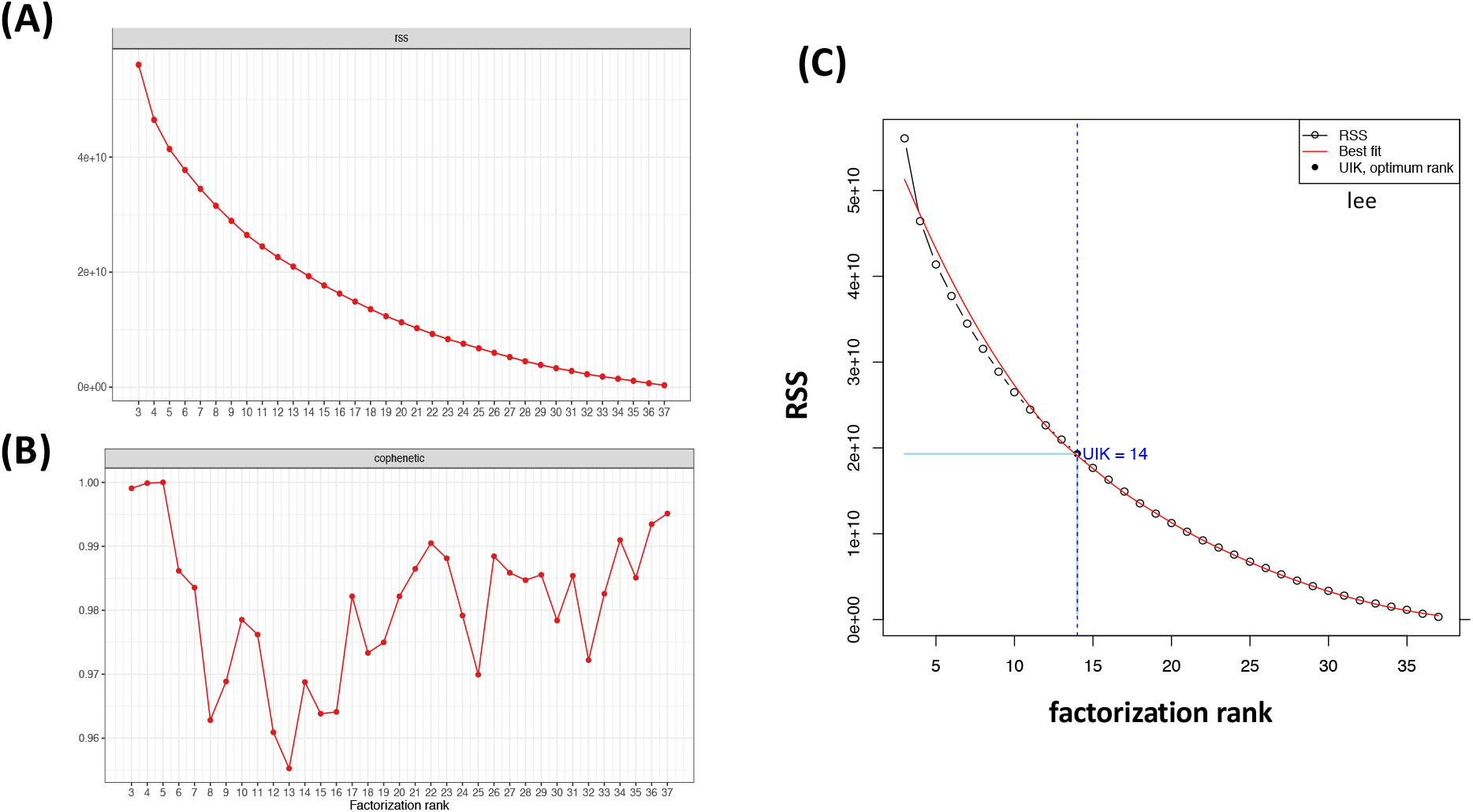
(A and B) Rank survey plots for RSS and cophenetic coefficient curves factorization rank. Factorization rank ranges from 3 to 37. The aim is to decide the optimal rank factorization is very rigid by a sudden look. (C) The function of factorization rank selected as emergence rank of RSS survey. The rank range between knee points is detected by the uik() function of R package inflection at the curve of the cumulative rank units. The best fit is performed using a linear regression model.

The most standard approach for such a judgment is by taking the so-called ‘elbow point,’ which is virtually the point where a severely decreasing or increasing curve begins to turn ‘flat enough’ (12,13,16–18). Thus, this study considered the function of the rank factorization curve and used the function uik() from the R package *inflection* to select the optimal rank (17,19,20). The uik() function detects the factorization rank when the curve begins to climb faster (start point) and the point beyond which the curve flattens out (ending point), which are generally known as the *knee points* of a curve (Figure 1). Figure 1 selected as the emergence of factorization rank of Golub gene expression data set on the rank survey plot. The optimal rank of the RSS plot is in between *knee points* detected by the uik() function of R package *inflection* at the curve of the cumulative *rank factorization belongs*.

We note that various techniques have been developed to select the optimal rank factorization, such as Brunet et al., 2004 suggested seizing the first value of r for which the cophenetic coefficient value was decreasing (7). Frigyesi et al., 2008 considered the smallest value at which the decrease in the RSS is lower than the decay of the RSS simulated from random data (9).

### 2.4. Applications

#### 2.4.1. Experimental Data and Analysis Codes

Analyses were performed utilizing the R programming language. Before the analyses procedure, the low-quality genes with the inadequate number of reads were eliminated, and gene expression converted logarithmic scale. Then, the data set was normalized by computing the averages of each sample in R. This study uses the NMF package of R to draw plots of rank surveys using plot() function (21). Rank survey analysis was performed to compare optimal rank with distinct methods using inflection package’s uik() and check_curve() functions (19). To import the simulated MATLAB file into the R environment, we utilized the readMat() function of the R.matlab package (22).

#### 2.4.2. Gene Expression data set

This study illustrates the utilization of NMF based on the UIK method to select optimal rank on RSS curve with **leukemia (esGolub) gene expression data set** in simplifying cancer subtypes (23–25). It was used in several papers on NMF and was built in the NMF package’s data (7,14,26), packed into an ExpressionSet object. To achieve biologically meaningful results, we used the entire gene expression data that has 5000 features by 38 leukemia samples. The difference between acute myelogenous leukemia (AML) and acute lymphoblastic leukemia (ALL) has been noted. ALL is also separated into two subtypes that T and B cells. It is remarked that this data set has been a touchstone in the categorization task of cancer.

Furthermore, it is remarked that this data set has been a touchstone in cancer classification by molecul, histology, and stage (23,27). This study reprocesses this data set to compare several clustering techniques regarding their effectiveness and permanence in recuperating other differentially expressed genes and associated pathways. Before the NMF procedure, dimension reduction is recommended for the larger gene expression dataset by non-specific criteria based on the characteristics of the expression estimates, i.e., the mean threshold of variance and genes with the smallest average variances (28).

For example, by looking at the NMF rank survey plot of RSS in Figure.1, we want to decide how many basis vectors we should keep to obtain the optimal rank of the target (original) matrix. To achieve such a task, an unbiased technique for deciding the number of clusters without eye involvement simultaneously capable of utilizing a computational program is a need.

#### 2.4.3. Simulated Mutational Processes Data

The simulated mutational process data obtained from Alexandrov et al. and his coworkers (29) is publicly available as a MATLAB file on SigProfiler (30). They identified the handful of processes functional a group of 100 simulated cancer genomes founded on the repeatability of their signatures and low error for reconstructing the novel catalogs. They generated the data by employing ten mutational processes with different signatures (motifs), each with 96 mutation types, and adding a Poisson noise. The data also corresponds to the six substitution types and their immediate 5′ and 3′ sequence background.

## 3. RESULTS

### 3.1. An application of NMF based on UIK method: Leukemia (esGolub) Data Set

The results shown in this paper utilize the NMF package of Gaujoux (2010) combined with Hutchins (2008) technique (Figure 1), but as shown in Figure 2, this study also tested other algorithms taken from “brunet” and “nsNMF” methods to see remarkable differences (7,21). It is important to emphasize that there is no remarkable base in the experi-mental data we examine. Consequently, we cannot show further considerable doubt that our approach operates effectively on the experimental data set. In Figure 2, the uik() function selects the optimal rank as the curve starts to declines faster (start point) and the point beyond that the curve flattens out (ending point), which are generally known as the knee points of a curve (Figure 1).

**Figure 2.**
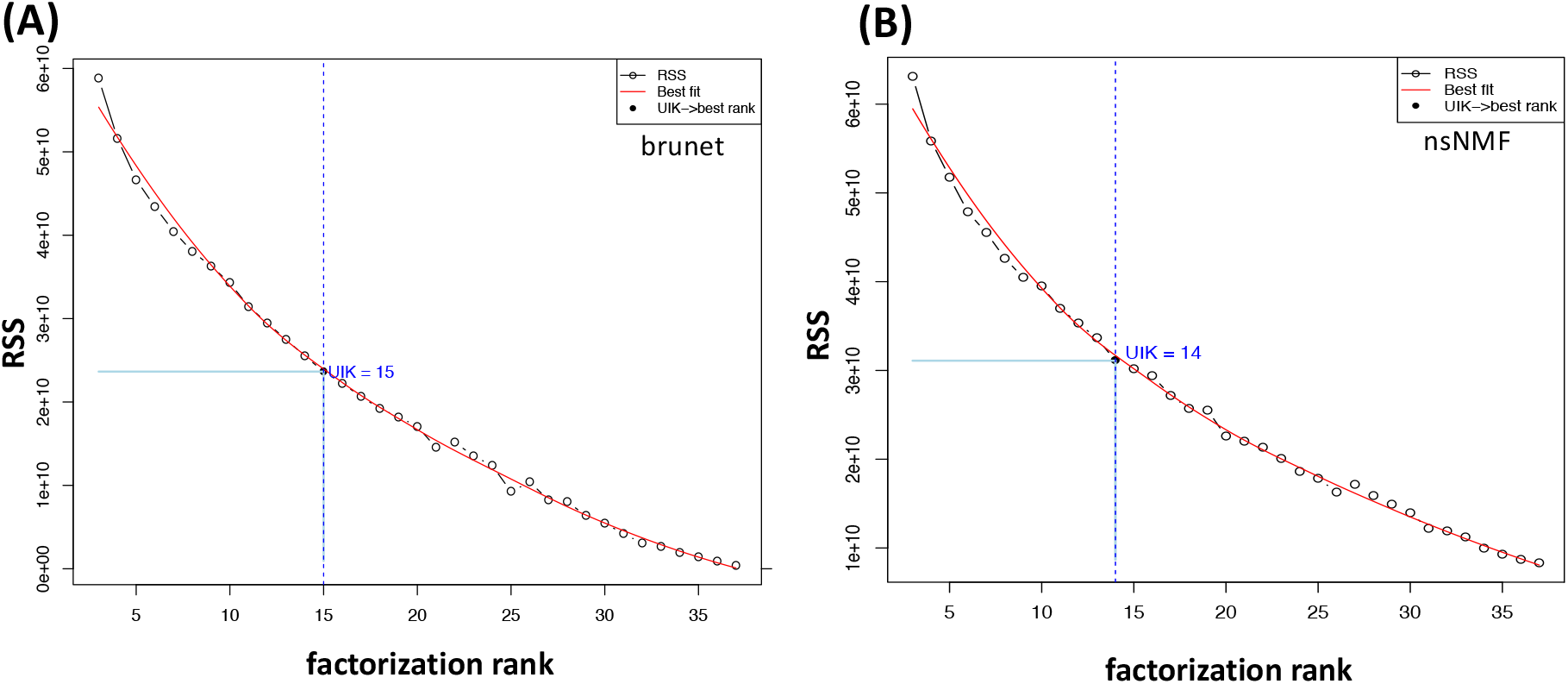
Application of UIK method on different algorithms (A) “brunet” and (B) “nsNMF”. The optimal rank, which UIK represents, is 15 for the brunet algorithm, whereas the UIK of nsNMF algorithm reveals the number 14 as an optimum rank likewise “lee” method.

By simply looking at the cophenetic correlation or RSS plots of rank factorization in Figure 3(A), one can confirm the optimum rank factorization is 3. For performance reasons, submatrix esGolub[1:200,] was previously performed here with only ten runs for each rank value. As demonstrated in Figure 3(B), we validated the UIK method of optimal rank factorization by comparing Gaujoux’s estimates of the esGolub sub-data set (31).

**Figure 3.**
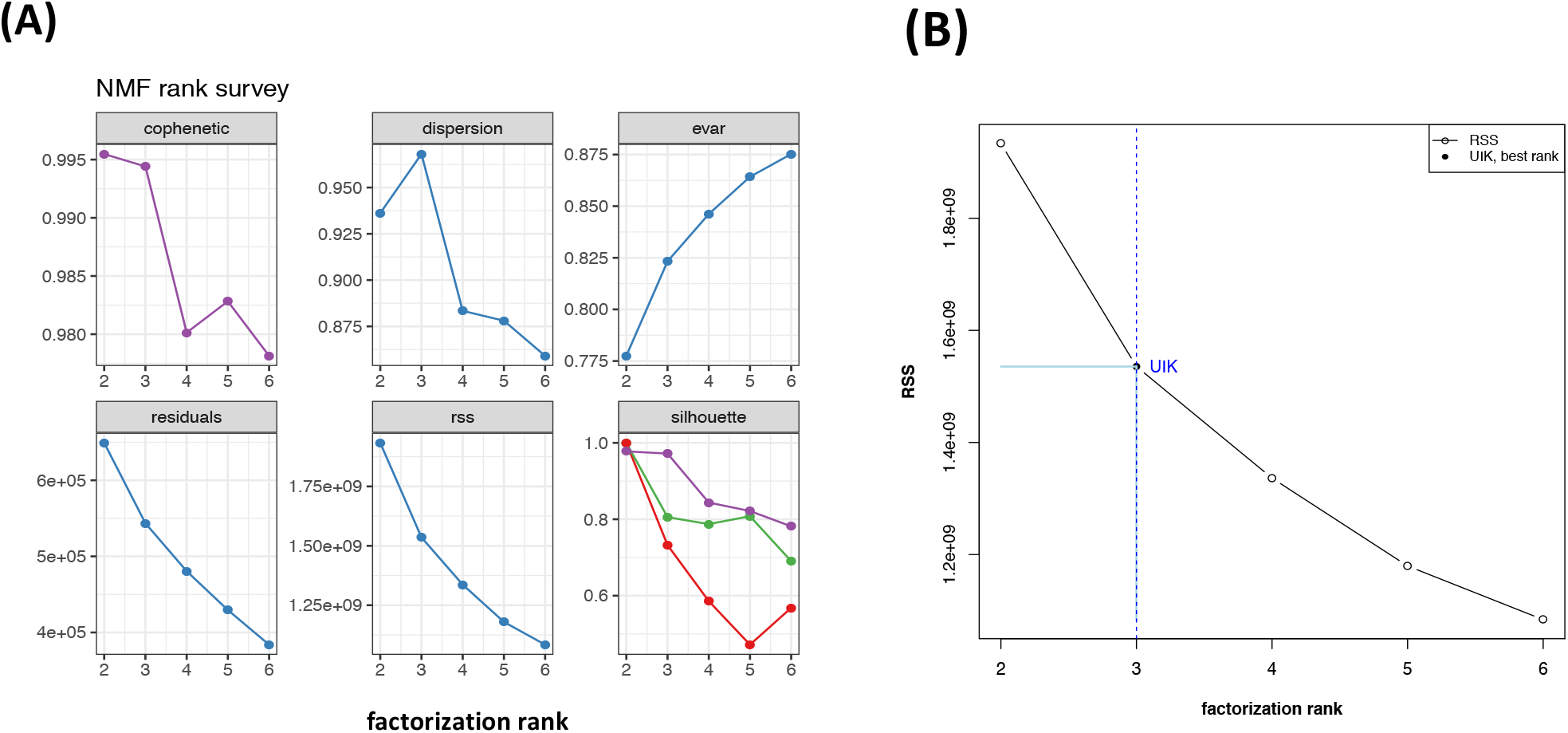
(A) Estimation of the optimal rank. NMF survey plot of quality measures obtained from factorization rank from 2 to 6 by running ten times of the target matrix esGolub[1:200,]. (B) The function of factorization rank is selected as the emergence rank of the RSS survey. For example, rank range 2 to 6 is between knee points detected by the R package inflection’s `uik()`function at 3. Overall, we confirmed the method of the UIK estimation with former results.

### 3.2 Application of NMF based on the UIK method: Simulated mutational process data

It is challenging to observe the rank factorization of the simulated data on the cophenetic coefficient curve (Figure 4A). Moreover, there is no clue deciding rank factorization based on Figure 4 just by observing the cophenetic correlation (Figure 4A) and the RSS (Figure 4B) plots. Nevertheless, the UIK method successfully validated their work and calculated ten mutational signatures for the simulated data.

**Figure 4.**
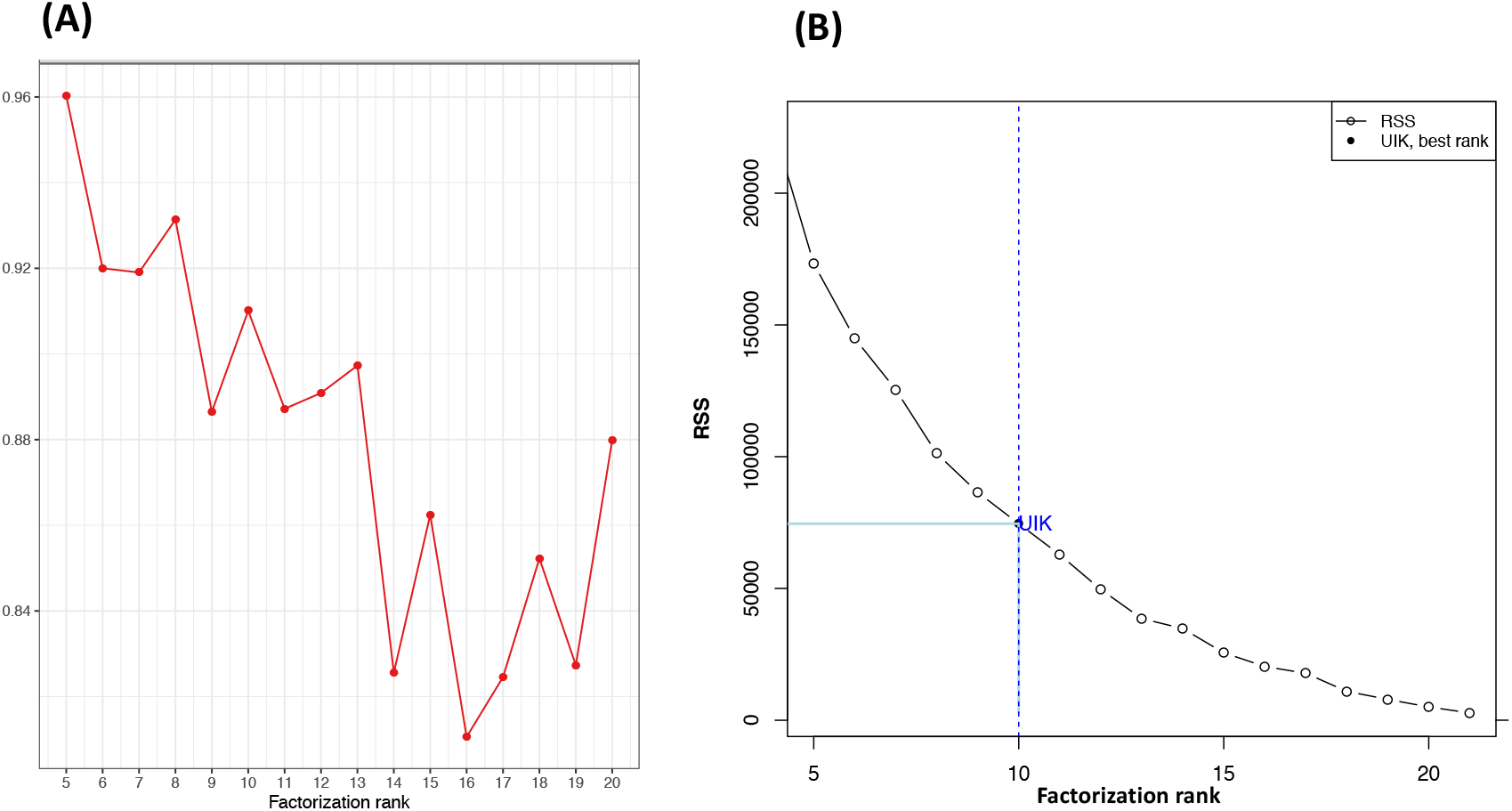
(A) It is complicated to locate the optimal rank with Brunet et al., 2008 approach. (B) However, the UIK method decides it faster and accurately, which agrees with the number of signatures detected by Alexandrov et al., 2013

## 4. CONCLUSIONS

Our novel finding is to apply the UIK method to selecting optimal rank based on the RSS curve of factorization ranks. First, this study employed Golub data set (23) and simulated data (29) utilizing UIK, which does not require average out the results from different runs of nmf() function or considering the variance between each run. In the second module, UIK precisely estimates simulated data with known dimensions. UIK technique tested on the Golub data set is likely to work both experimental data and synthetic or simulation data. Moreover, finally, the technique of UIK is free of a priori rank parameter input and does not require setting initial parameters that considerably affect the performance.

This study shows the optimal rank of an NMF model for gene expression or mutational processes datasets utilizing the UIK method. The UIK method tested on gene expression data deconvolution can achieve optimal rank estimation. This study also shows that the UIK method provides a credible prediction for gene expression data and precisely estimates simulated data with known dimensions. Our proposed UIK method based on RSS curvature first inflection point to estimate optimal rank is theoretically superior or equivalent to existing implementation and software. All the undertaking is done with R programming and is freely available.

The analysis has some limitations, such that other NMF packages or software on the gene expression research are not tested. The study demonstrates that the UIK method provides a credible prediction for gene expression data and However, simply assuming they used the same algorithms of NMF, as far as RSS and residual curves would be approximated the same, the UIK method would result in the same optimal ranks.

## Supporting information

https://www.dropbox.com/s/5dz6lbo1n7dv761/SUPPLEMANTARY%20FILE%20S4_uikmethod.pdf?dl=0

## HUMAN AND ANIMAL RIGHTS

Not applicable

## CONSENT FOR PUBLICATION

Not applicable.

## ACKNOWLEDGEMENTS

Emine Güven designed the project, performed the computations, analyzed the data, contributed reagents/materials/analysis tools, wrote the paper, prepared figures and/or tables, reviewed drafts of the paper.

## SUPPORTIVE/SUPPLEMENTARY MATERIAL

The data supporting the findings of the article is available at <u>Supplementary Files</u>.

