## Supplementary material for "The Decision of the Optimal Rank of a Non-negative Matrix Factorization Model for Gene Expression Datasets Utilizing Unit Invariant Knee Method": https://www.dropbox.com/s/5dz6lbo1n7dv761/SUPPLEMANTARY%20FILE%20S4_uikmethod.pdf?dl=0

SUPPLEMANTARY FILES

Emine Güven[a]

[a]*Biomedical Engineering Department, Faculty of Engineering, Düzce University, Düzce, Turkey*

#### Supplementary File S1

Alexandorav et al., 2013 the simulated mutational signatures data summary

| | | |
|---|---|---|
| simulated_data | list [5] | List of length 5 |
| cancerType | character [1 x 1] | 'WTSI BRCA Genomes' |
| originalGenomes | double [96 x 21] | 31 34 9 21 13 15 110 91 9 87 100 46 122 112 13 107 52 ... |
| sampleNames | list [21] | List of length 21 |
| subtypes | list [96] | List of length 96 |
| types | list [96] | List of length 96 |

Supplementary File S2

**Implementation of the comparison of Gaujoux's estimates of the esGolub sub-data set as with the UIK method**

```
require(NMF)
require(inflection)

data("esGolub")
sub_esGolub<-esGolub[1:200,]
estim.r<-nmfEstimateRank(sub_esGolub,r=2:6,nrun=10,seed=123456)
rankSurvey<-estim.r

pdf("plots/repeatEsgolub.pdf")
plot(rankSurvey)
dev.off()

x=rankSurvey$measures$rank
y=rankSurvey$measures$rss

rank=uik(x,y)

x1=min(x)
x2=max(x)
pdf("plots/uik_repeatEsgolub.pdf")
plot(x,y,type="b",xlab="factorization rank",ylab="RSS", pch=c(1,16,1,1,1),
                  xlim=c(2,6),font.lab=2,cex=1)
segments(rank,y[2],rank,y[5], lwd=2, pch=16,col="lightblue")
segments(x[1],y[2],x[2],y[2], lwd=2,pch=16, col="lightblue")

abline(v=rank,col='blue',lty=2)
axis(side=4,at=rank,labels=as.character(rank),cex.axis=0.6)
text(x = rank+0.2, y[2], label = "UIK",col="blue")
legend("topright",c("RSS","UIK, best rank"),cex=.8,
       col=c("black","black"),lty=c(1,NA),pch=c(1,16))

dev.off()
```

Supplemantary File S3

**The Rank survey plot of Alexandorav et al., 2013 simulated mutational signatures data**

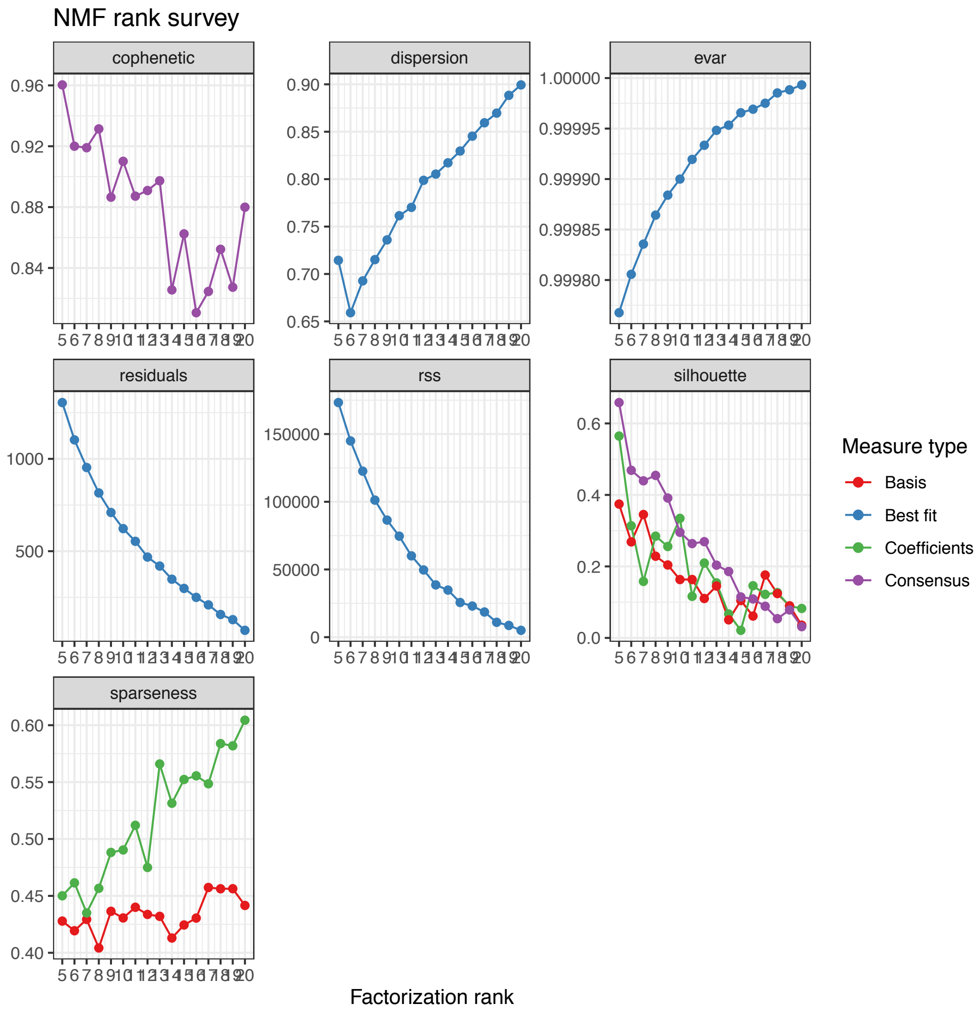

### Supplementary File S4

**The UIK method on Alexandorav et al., 2013 simulated mutational signatures data**

```
library(NMF)
```

```
## Loading required package: pkgmaker
## Loading required package: registry
## Loading required package: rngtools
## Loading required package: cluster
```

Text

```
## NMF - BioConductor layer [OK] | Shared memory capabilities [OK] | Cores 3/4
```

```
library(inflection)
library(R.matlab)
```

```
## R.matlab v3.6.2 (2018-09-26) successfully loaded. See ?R.matlab for help.
##
## Attaching package: 'R.matlab'
## The following objects are masked from 'package:base':
##
##     getOption, isOpen
```

```
data <- readMat('21_breast_WGS_substitutions.mat')
summary(data)
```

```
##                 Length Class  Mode
## cancerType         1   -none- character
## originalGenomes 2016   -none- numeric
## sampleNames       21   -none- list
## subtypes          96   -none- list
## types             96   -none- list
```

```
str(data)
```

```
## List of 5
##  $ cancerType     : chr [1, 1] "WTSI BRCA Genomes"
##  $ originalGenomes: num [1:96, 1:21] 31 34 9 21 13 15 8 11 41 17 ...
##  $ sampleNames    :List of 21
##   ..$ :List of 1
##   .. ..$ : chr [1, 1] "PD3851a"
##   ..$ :List of 1
##   .. ..$ : chr [1, 1] "PD3890a"
##   ..$ :List of 1
##   .. ..$ : chr [1, 1] "PD3904a"
##   ..$ :List of 1
##   .. ..$ : chr [1, 1] "PD3905a"
##   ..$ :List of 1
##   .. ..$ : chr [1, 1] "PD3945a"
##   ..$ :List of 1
##   .. ..$ : chr [1, 1] "PD4005a"
##   ..$ :List of 1
##   .. ..$ : chr [1, 1] "PD4006a"
##   ..$ :List of 1
##   .. ..$ : chr [1, 1] "PD4085a"
##   ..$ :List of 1
##   .. ..$ : chr [1, 1] "PD4086a"
##   ..$ :List of 1
```

```
##    .. ..$ : chr [1, 1] "C>T"
##    ..$ :List of 1
##    .. ..$ : chr [1, 1] "C>T"
##    ..$ :List of 1
##    .. ..$ : chr [1, 1] "C>T"
##    ..$ :List of 1
##    .. ..$ : chr [1, 1] "C>T"
##    ..$ :List of 1
##    .. ..$ : chr [1, 1] "T>A"
##    ..$ :List of 1
##    .. ..$ : chr [1, 1] "T>A"
##    ..$ :List of 1
##    .. ..$ : chr [1, 1] "T>A"
##    ..$ :List of 1
##    .. ..$ : chr [1, 1] "T>A"
##    ..$ :List of 1
##    .. ..$ : chr [1, 1] "T>C"
##    ..$ :List of 1
##    .. ..$ : chr [1, 1] "T>C"
##    ..$ :List of 1
##    .. ..$ : chr [1, 1] "T>C"
##    ..$ :List of 1
##    .. ..$ : chr [1, 1] "T>C"
##    ..$ :List of 1
##    .. ..$ : chr [1, 1] "T>G"
##    ..$ :List of 1
##    .. ..$ : chr [1, 1] "T>G"
##    ..$ :List of 1
##    .. ..$ : chr [1, 1] "T>G"
##    ..$ :List of 1
##    .. ..$ : chr [1, 1] "T>G"
##  - attr(*, "header")=List of 3
##    ..$ description: chr "MATLAB 5.0 MAT-file, Platform: MACI64, Created on: Thu Feb 15 20:52:02 2018
##    ..$ version    : chr "5"
##    ..$ endian     : chr "little"
```

```r
originalGenome<-data$originalGenomes
head(originalGenome)
```

```
##      [,1] [,2] [,3] [,4] [,5] [,6] [,7] [,8] [,9] [,10] [,11] [,12] [,13] [,14]
## [1,]   31  110  122   94  243   74  198   61   31    34   112   210   122   228
## [2,]   34   91  112   69  163   66  173   51   19    18    50   176   133   169
## [3,]    9    9   13   11   24   12   33    7    8     3     7    33    12    21
## [4,]   21   87  107   65  155   64  192   49   23    26    44   176    96   158
## [5,]   13  100   52   66  130   56  164   18   18    15    30   126    95   128
## [6,]   15   46   42   41   78   34   96   18   14    11    28    85    72    74
##      [,15] [,16] [,17] [,18] [,19] [,20] [,21]
## [1,]   128   165    48    28    58    58    64
## [2,]   138    55    42    24    72    36    43
## [3,]    15    20    16     6    12    13     7
## [4,]   142    65    40    13    45    37    35
## [5,]   102   191    60     8    37    23    25
## [6,]    71    72    26     5    27    19    18
```

```
### nmf rank suvey
simulated.r<-nmfEstimateRank(originalGenome,r=2:21,nrun=35,seed=123456)

#simulated.r2<-nmfEstimateRank(originalGenome,r=5:20,nrun=30,seed=123456)
rankSurvey<-simulated.r
#rankSurvey<-simulated.r2

#pdf("plots/repeatEsgolub.pdf")
plot(rankSurvey)
```

```
## Warning: Removed 1 row(s) containing missing values (geom_path).

## Warning: Removed 1 rows containing missing values (geom_point).
```

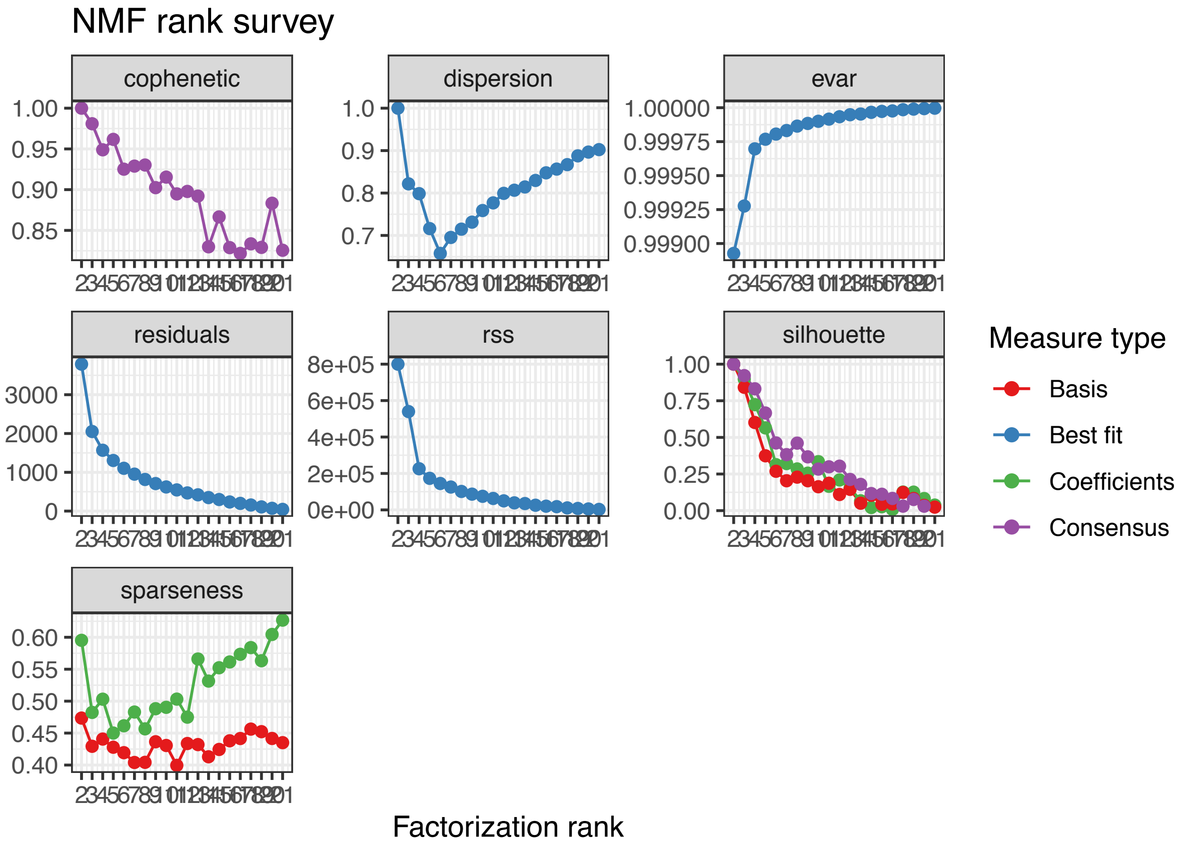


```
#dev.off()
```

```
x=rankSurvey$measures$rank
y=rankSurvey$measures$rss

rank=uik(x,y)



#plot(x,y,pch=19,cex=0.9)
x1=min(x)
x2=max(x)
#pdf("plots/uik_repeatEsgolub.pdf")
plot(x,y,type="b",xlab="factorization rank",ylab="RSS",pch=c(rep(1,8),16,rep(1,11)),xlim=c(5,21),font.la
```

```r
#par(new=TRUE)
#plot(rank,y[4],pch=16,col="blue", xaxt='n', yaxt='n',ann=FALSE)
segments(rank,y[9],rank,y[20], lwd=2, pch=16,col="lightblue")
segments(x[1],y[9],x[9],y[9], lwd=2,pch=16, col="lightblue")

abline(v=rank,col='blue',lty=2)
axis(side=4,at=rank,labels=as.character(rank),cex.axis=0.6)
text(x = rank+0.4, y[7], label = "UIK",col="blue")
#srt = 90,col='navyblue') # Rotation

legend("topright",c("RSS","UIK, best rank"),cex=.8,
       col=c("black","black"),lty=c(1,NA),pch=c(1,16))
```

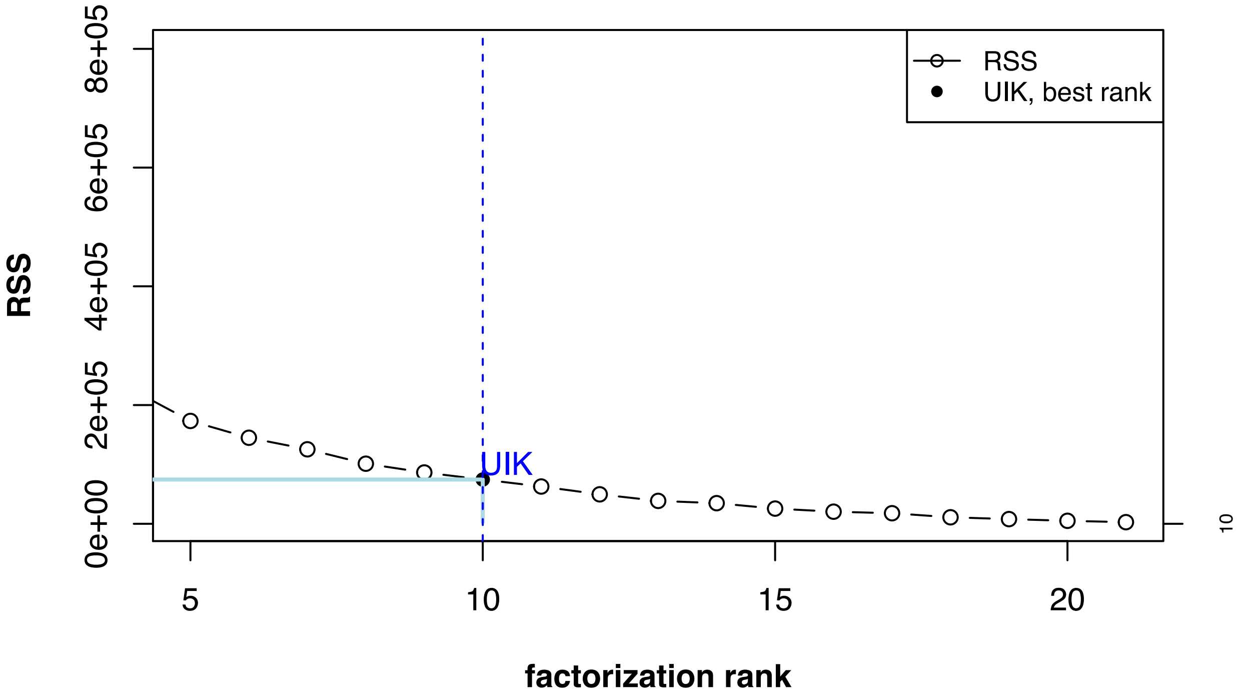


```r
#dev.off()
```
